# scDAPP: a comprehensive single-cell transcriptomics analysis pipeline optimized for cross-group comparison

**DOI:** 10.1101/2024.05.06.592708

**Authors:** Alexander Ferrena, Xiang Yu Zheng, Kevyn Jackson, Bang Hoang, Bernice Morrow, Deyou Zheng

## Abstract

Single-cell transcriptomics profiling has increasingly been used to evaluate cross-group differences in cell population and cell-type gene expression. This often leads to large datasets with complex experimental designs that need advanced comparative analysis. Concurrently, bioinformatics software and analytic approaches also become more diverse and constantly undergo improvement. Thus, there is an increased need for automated and standardized data processing and analysis pipelines, which should be efficient and flexible too. To address these, we develop the **s**ingle-**c**ell **D**ifferential **A**nalysis and **P**rocessing **P**ipeline (scDAPP), a R-based workflow for comparative analysis of single cell (or nucleus) transcriptomic data between two or more groups and at the levels of single cells or “pseudobulking” samples. The pipeline automates many steps of pre-processing using data-learnt parameters, uses previously benchmarked software, and generates comprehensive intermediate data and final results that are valuable for both beginners and experts of scRNA-seq analysis. Moreover, the analytic reports, augmented by extensive data visualization, increase the transparency of computational analysis and parameter choices, while facilitate users to go seamlessly from raw data to biological interpretation.

**Availability and Implementation:** scDAPP is freely available for non-commercial usage as an R package under the MIT license. Source code, documentation and sample data are available at the GitHub (https://github.com/bioinfoDZ/scDAPP).

## Introduction

Advancements in single-cell transcriptomics technologies and reductions in cost have greatly increased the scale and complexity of experiments using single-cell RNA-seq (scRNA-seq) and single-nucleus RNA-seq (snRNA-seq)^1^. In 2023, nearly 4,000 papers were published using scRNA-seq, illustrating an exponential increase over the past 5 years (PubMed). Not only have these methods been used for simple categorization of cell population in tissues or biological samples, but also increasingly as read-outs of population and gene program changes across experimental conditions, such as genetic knockouts or drug treatments. As the technologies for data acquisition become more sophisticated, the bottleneck of innovation has shifted to efficient and rigorous bioinformatic analysis, to guide investigators to assess data quality rapidly and use benchmarked software for uncovering biological signals efficiently and robustly. Standard, scalable and modular workflows would greatly facilitate this process.

As single cell data acquisition technologies advance, bioinformatic algorithms and software for data analysis have also evolved rapidly and become challenging for ordinary researchers to follow. Various benchmark comparisons of software, however, have provided excellent recommendations for selecting methods for most data analysis steps, such as statistically rigorous strategies for cross-group comparisons, including differential expression analysis and differential cell composition analysis^2–5^. Such studies have shown that the most important criterion for robust methodological performance is the capacity to explicitly model biological replicate variability via “pseudobulking”. It was demonstrated that the alternative of treating data of individual cells as independent measurements could easily lead to inflated and inappropriate statistics, resulting in less biologically relevant and small effect-size discovery that is prone to noises. Related to this, advancements in sample multiplexing, such as Cell Hashing, Multi-Seq, or 10X Cell Multiplexing, have significantly increased the number of replicates (at reduced cost) and complexity of data analysis^6,7^. Likewise, our previous study demonstrated that the Reference Principal Component Integration (RPCI) algorithm, released in the Robust Integration of Single Cell (RISC) RNA-seq software package, could integrate multiple-sample data with high accuracy, while avoiding over-correction, and with the ability to easily return batch-corrected expression values^8^.

We present here an R-based pipeline called **s**ingle-**c**ell **D**ifferential **A**nalysis and **P**rocessing **P**ipeline (scDAPP) for cross-group comparative analysis of scRNA-seq and snRNA-seq data from 10X Genomics platform. Our design emphasizes scalability, ease of use, user-friendly graphic visualization, transparency, and reproducibility. Similar pipelines exist, including Cellsnake, Cellenics, scDrake, SingleCAnalyzer, and the Single-Cell Omics workbench^9–13^. Compared to them, scDAPP focuses more on comparative analysis across groups. As such, it implements critical options for using replicates to generate pseudobulk data automatically, which are more appropriate for cross-group comparisons, for both gene expression and cell composition analysis, as discussed above. Furthermore, it supports complex multi-group comparisons (such as Drug A vs Drug B vs Control) and comprehensive downstream bioinformatics analysis, such as Gene Set Enrichment Analysis (GSEA) and transcription factor target analysis. Additionally, flexible input formats allow for direct data importing from raw CellRanger outputs or pre-processed scRNA-seq data objects. Importantly, using enriched visualization, scDAPP is designed to guide users to examine and understand the selection of parameters in each step of the data analysis, so that users can have a good grasp of the options and make appropriate and rational adjustments. The enriched visualization is especially valuable because it shows the underlying data distributions and moreover how parameter choices affect the analytic results. Overall, scDAPP facilitates and systematizes data processing steps and allows users to quickly delve into biological interpretation, thus advancing the rigor and scalability of single-cell transcriptomic data analysis.

### Overview of scDAPP: Core features and run configuration

The scDAPP wraps previously published software packages into R markdown codes, with critical pipeline-specific implementation (**Figure 1)**. It starts with several cell filtering functions using parameters learnt from input data’s distribution, including doublet removal, followed by individual sample analysis, integration of samples, clustering of the integrated data, and cross-group comparisons of cell cluster abundance and gene expression using either cell-level or pseudobulk-level data (when replicates are used). The core packages include Seurat for individual sample analysis and data visualization (v5 and up are supported), RISC for sample integration and batch correction, the Speckle package with the Propeller test for replicate-aware (“pseudobulk”) cross-condition cell composition comparison, EdgeR for pseudobulk-level differential gene expression analysis across conditions, and FGSEA for pathway analysis^8,14–17^. Importantly, multigroup comparative analysis is supported (i.e., A vs B vs C). Additionally, label transfer from previously cell-type annotated data is an optional feature to facilitate cell cluster annotation.

**Figure 1.**
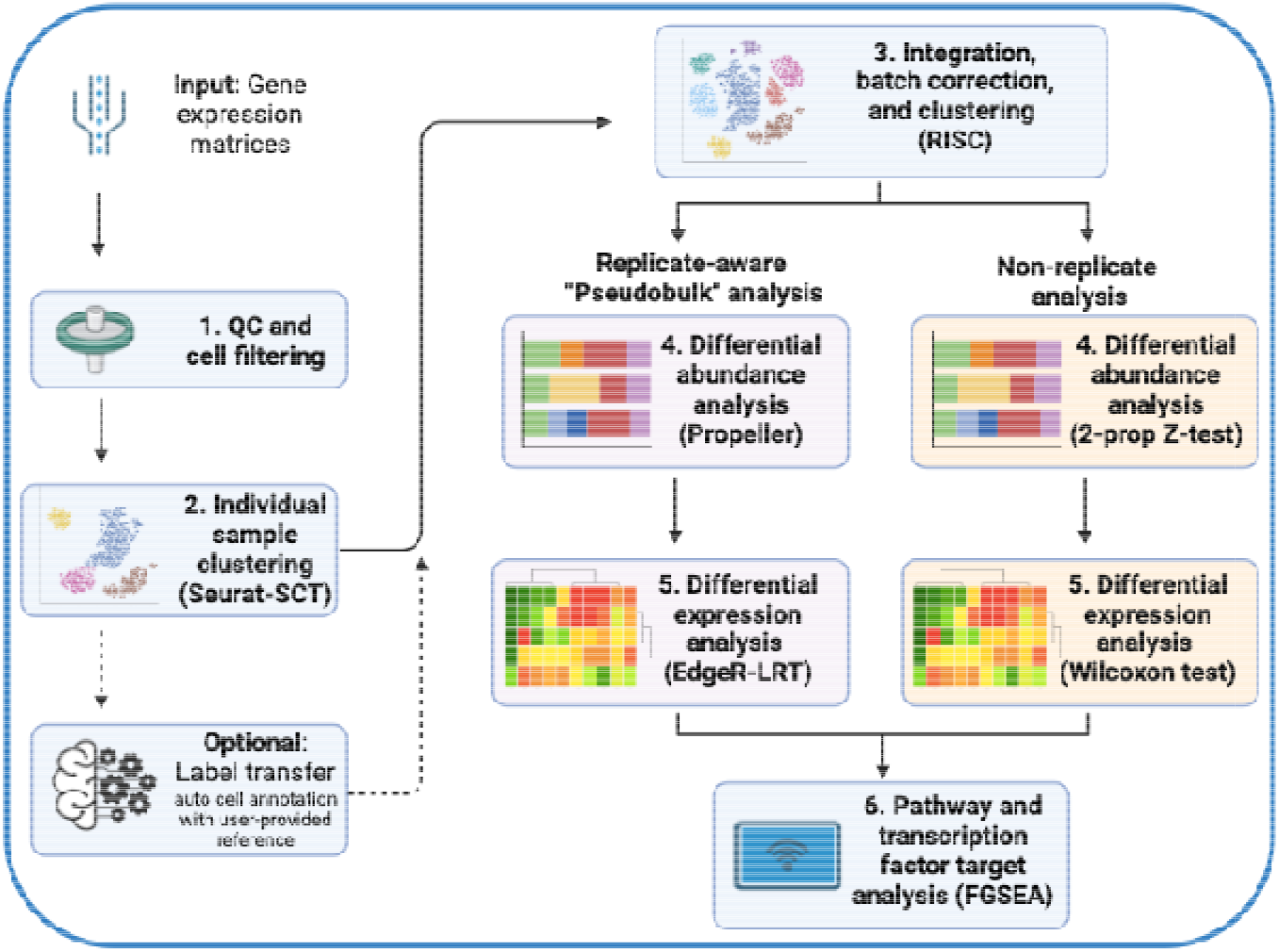
Scheme of scDAPP: main components and software.

Key input options and formats for the pipeline are shown in **Figure 2**. The input for scDAPP is the raw (non-normalized) UMI (Unique Molecular Identifiers) counts matrix for each sample. This can be either the filtered matrices from CellRanger (“filtered_feature_bc_matrix.h5” files) or Seurat objects. The latter option allows for flexibility, for example, to use data that have been filtered or processed by other means. In addition, pipeline run configuration files are needed, which specify sample-wise metadata including per-sample information and optional sample nickname codes (“sample_metadata”), and a list of the relevant cross-group comparisons (“comps”).

**Figure 2.**
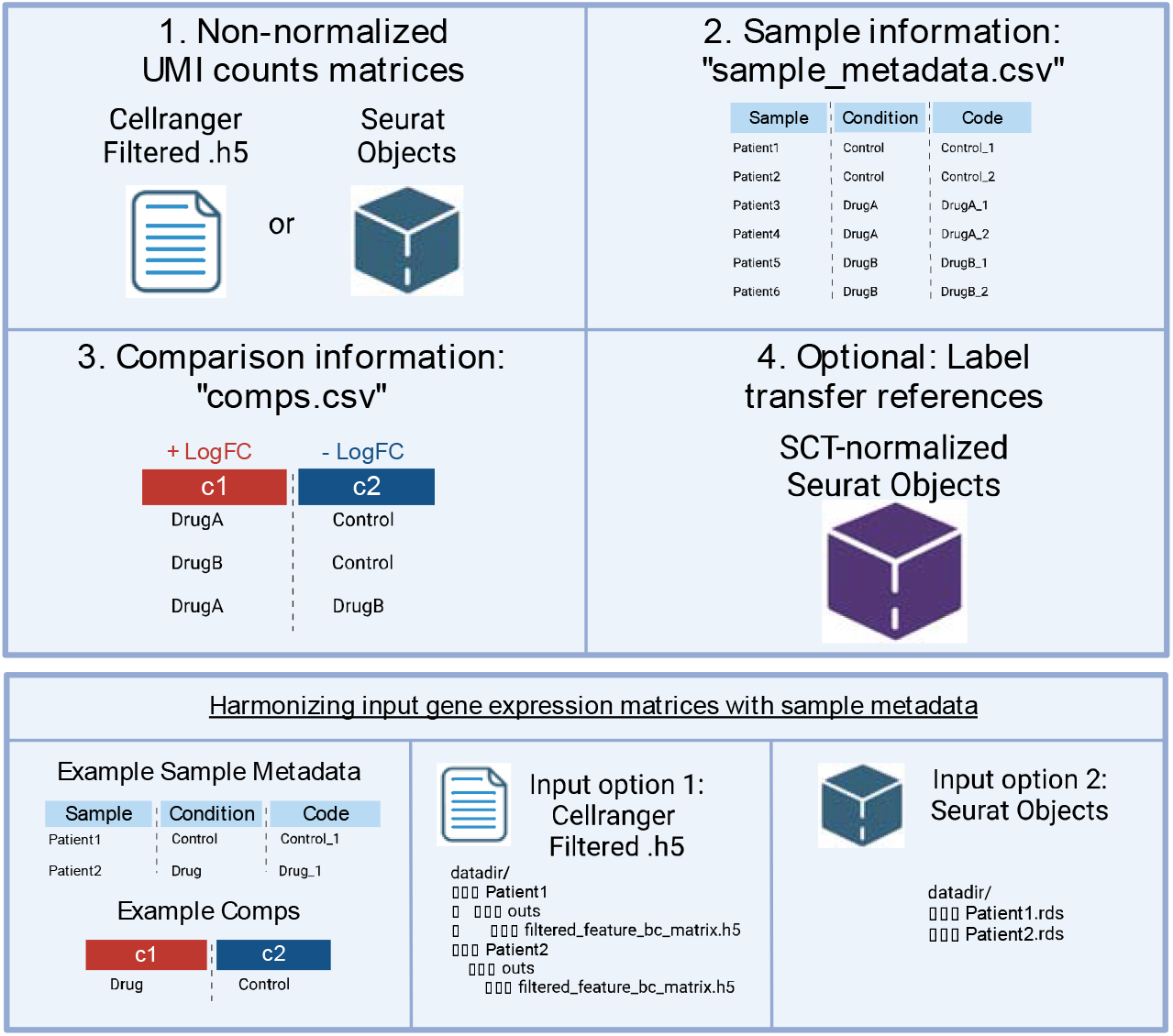
Inputs for scDAPP: data format and options.

One critical input parameter is “Pseudobulk_mode” (TRUE/FALSE) that will specify whether to make cross-group comparisons using samples in the same group/condition as replicates. Setting it to “TRUE” will run scDAPP in a replicate-aware manner, which is highly recommended when biological replicates are available. It has two effects: 1). Differential gene expression analysis will be run in a pseudobulk manner, invoking the EdgeR-Likelihood Ratio Test on the aggregated reads from all cells in each cluster in each sample. This was shown to perform more accurately than using data of each cell as an independent measurement (leading to inflated sample size and thus statistics) ^2^. 2). Differential cell composition analysis will also be run in a replicate-aware manner via the “Propeller” test from the “speckle” package, which was demonstrated to perform more accurately in a recent benchmark analysis^3^. Conversely, setting the “Pseudobulk_mode” parameter to “FALSE” will evoke scDAPP to perform differential gene expression analysis by the Wilcoxon test in Seurat at the single cell level (i.e., treating each cell as an independent data point), and differential cell abundance analysis by the two-proportion Z test as implemented in the R “prop.test” function. This non-replicate option is only recommended for comparative analysis without replicates but can be run on inputs with replicates. It is more prone to false positive, but this is sometimes unavoidable, such as in the context of pilot studies.

Two other key optional parameters are provided in the run configuration file. The first is “use_labeltransfer” for invoking the use of the Seurat label transfer workflow. If set to “TRUE”, two additional parameters need to be set: one refers to a Seurat object containing normalized data from the Seurat’s SingleCellTransform workflow and a metadata column called “Celltype” to be used for label transfer, and the other (“m_reference”) points to a file listing marker genes for the reference cell types in the output format of the Seurat “FindAllMarkers” function. The second key option is the “risc_reference” parameter, specifically related to the RISC software used for integration and batch correction. The RPCI algorithm within RISC uses one of the input samples to learn the principal component space to project cells in all samples. Users can specify which sample to be the RPCI reference, after they examine the results from all samples. Alternatively, and by default, scDAPP provides an automated RISC reference selection algorithm (described below).

The pipeline has additional fields related to quality control metric thresholds, tuning of data-driven cell filtering, and hyperparameter selection in clustering analysis, with reasonable defaults, as described below. One more required input is “species”, which is used to search the MSIGDB database for the correct gene symbols during pathway enrichment analysis, via the msigdbr package^18^.

### QC and cell filtering

After reading in the data, the first step in scDAPP is quality control (QC) and filtering of poor-quality cells. On top of pre-set lenient thresholds, scDAPP tries to learn better cutoffs from the input data. Currently scDAPP considers the following information for cell quality filtering: number of UMIs per cell, number of unique genes (i.e., features) per cell, percent of UMIs from the mitochondrial genes per cell, and the prediction of doublets and multiplets. Additionally, though not commonly implemented in other software, scDAPP evaluates the percent of reads from hemoglobin-related genes, as red blood cell lysis buffer may miss its target population; these cells have the very distinct feature of extremely high hemoglobin gene expression. For each of these QC metrics, users may select initial lenient thresholds, such as minimum UMIs (500 by default), minimum number of unique genes (200), maximum percent of mitochondrial reads (25%), and maximum percent of hemoglobin read (25%). On top of these users’ settings, scDAPP will further optimize the thresholds by analyzing the distribution of the underlying data. The distributions and the corresponding data-driven cutoffs are presented to the users graphically so that they may select thresholds deemed more appropriate. Essentially, the algorithms for deriving these data-driven cutoffs try to emulate the common manual QC, such as visual inspection of violin plots of these variables. First, a “complexity” filter is applied, removing cells with a lower-than-expected number of unique genes given the number of observed UMIs. This is determined by two regression models, linear regression and LOESS regression with the log(nGenes) as the dependent variable and log(nUMI) as the predictor. Cells are considered low-complexity outliers if their linear regression Cook’s distance is greater than 4 / number of cells, and their LOESS regression scaled residual value is less than −5, by default. The residual threshold is passed as a user parameter with higher values increasing the strictness of the filter. Next, scDAPP will learn data-driven cutoffs for the number of UMIs and the percent of mitochondrial reads (%mt). For these, robust statistics methods are applied, where cells with median absolute deviations above +2.5 (by default, tunable) for %mt and below −2.5 (default) for number of UMIs are considered low-quality outliers and removed. Again, users have the option to ignore these data-driven cutoffs entirely.

After poor quality cells are removed, scDAPP optionally uses DoubletFinder to predict doublets for removal from further analysis^19^. DoubletFinder hyperparameters such as the homotypic doublet rate are automatically estimated for each sample using the number of cells and the empirical multiplet rate provided by 10X Genomics^20^.

### Individual sample clustering analysis

After cell filtering, individual sample analysis is performed with Seurat using the modified SingleCellTransform (SCT) workflow^14,21^. For each sample, scDAPP applies SCT, principal component analysis (PCA), graph construction, Louvain clustering, and Uniform Manifold Approximation and Projection (UMAP) for visualization. The hyperparameters including number of PCs (30 by default) and Louvain resolution (default 0.5) can be specified by the user, and iterated if needed, after examining the result from default settings.

While cluster markers are always computed, cluster annotation by label transfer is an option, which uses the Seurat-based label transfer workflow and a reference Seurat object as described in the input section above. Two extensions are made in scDAPP to help users. 1). We apply a hard cutoff of label transfer score of 0.3, below which cells are considered non-classified, and a soft threshold of 0.5, below which cells are considered only putatively classified. 2). Seurat label transfer gives a score and label for each cell, but we extend this to the cluster level by setting the transferred annotation to the most common predicted cell type for each cluster. To show the quality of label transfer, scDAPP generates cluster-level violin plots for the label transfer scores and provides heatmap visualization of the expression of the reference marker genes. Automated cell calling tools like label transfer are useful for providing a suggested cell annotation, but the results should be carefully examined and thus scDAPP makes all the relevant plots and scores available to the users.

### Sample integration and batch correction

After individual samples are clustered, scDAPP uses the RISC workflow for integration and batch correction, starting from the raw data before Seurat analysis. RISC is also used for Louvain clustering and UMAP visualization of the integration data. There are scDAPP-specific extensions of the RISC package. By default, RISC uses the intersect of genes from the single cell objects as integration features, but this can leave out genes detected in only one sample. This can be problematic if one cell type (or corresponding marker genes) is absent in one sample or one of the comparison groups, potentially leading to missing marker genes for that cell type after integration. To overcome this, after cell filtering and clustering of individual samples, scDAPP reads the raw count data directly to RISC, selects cells used in the individual sample analysis, concatenates the cell-filtered matrices, and then performs gene filtering, such that more genes are retained. Several other key parameters are allowed for RISC integration, including the number of PCs, the number of neighbors during graph construction, and clustering resolution. The default parameters for these in scDAPP are reasonable: PCs = 30, resolution = 0.5, neighbors = 10, but they may be modified by users.

Another important extension of the RISC package in scDAPP is automated selection of a reference sample for multiple-sample integration. Currently, this is done manually. RISC users examine a panel of plots generated by the “InPlot” function of RISC, which describe the number of clusters, variance per PC, and a measure of distributional divergence for each sample ^8^, and then select a sample for integration reference. We decided to automate this process in scDAPP by calculating a heuristic reference score for each sample. This score is based on the number of clusters, the cluster diversity, and the number of cells in each sample, and generally the sample with the greatest number of clusters, greatest diversity of clusters, and highest number of cells gets the highest score and is chosen as the best reference.

For each sample *i*, let *n*_*i*_ denote the number of clusters, and *c*_*i*_ denote a weighted cell number defined as: 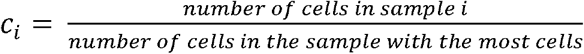

Next, let *v*_*ij*_ denote the variance of cluster *j* in sample *i*. We compute a cluster-average variance score *s*_i_ for sample *i* by calculating the mean value of the variances of each cluster:

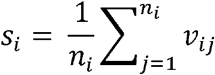

Then, a reference score *S*_i_ for sample *i* is computed as:

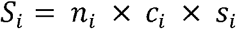

Finally, the sample *i*^*^ with the maximum *S* score is chosen as the reference: *i*= arg max_*i*_ *S*_*i*_.

This is meant to automate the RISC recommendation that the sample with the most cell types is the preferred integration reference. As with the setting of all other parameters in scDAPP, the pipeline shows all the RISC InPlot panels so that users can examine the underlying data and manually specify the reference sample via the “risc_reference” parameter in the run configuration file. Additionally, scDAPP generates alluvial plots to illustrate cluster relationships between individual samples and integrated data, thus helping users to spot over- or under-integration.

### Cross-group comparison

After RISC integration, scDAPP performs cross-group comparison. First, differential cell composition analysis is applied to study cell population changes between groups by comparing the proportions of cells in each cluster across samples. As mentioned above, this can be done in a replicate-aware manner using the “Propeller” test from the “Speckle” package, by setting the parameter “Pseudobulk_mode” = “TRUE”. In short, this is a t-test of the proportions for each cluster with samples in each group as replicates. Notably, scDAPP applies the square-root arcsine transformation that is provided as an option by Propeller, as this transformation was shown to perform best in a benchmarking analysis^3^. If replicates are not available or not considered, scDAPP will run in a non-replicate aware manner by setting “Pseudobulk_mode” = “FALSE”. This will utilize the two proportion Z-test as implemented in the R prop.test function.

Next, cross-group differential gene expression analysis is performed. As stated above, this can be done via a replicate-aware pseudobulk based method by setting “Pseudobulk_mode” = “TRUE”. In this scheme, the EdgeR likelihood ratio test (EdgeR-LRT) method is used, as this slightly outperformed default EdgeR and other pseudobulk methods like DESeq2 in a recent benchmark^2^. The data used for EdgeR are raw counts (i.e., UMIs) aggregated over cells in each cluster per sample for individual genes. The EdgeR statistical output is combined with other important single-cell information including the percent of cells expressing a gene in each condition, allowing downstream prioritization or further filtering of the differentially expressed genes (DEGs). If replicates are not available or not considered, differential expression can be run with “Pseudobulk_mode” = “FALSE” and performed using the Wilcoxon Rank-Sum test as implemented in the Seurat “FindMarkers” function, but importantly using the RISC batch-corrected gene expression values.

Finally, scDAPP performs multiple types of function enrichment analysis using the differential gene expression results. This includes Gene Set Enrichment Analysis (GSEA) as implemented in the “fgsea” R package^22^. Notably, GSEA allows pre-ranking of all genes based on differential expression summary statistics. This obviates the need for arbitrary differential expression thresholds and works well with pseudobulk-based methods. Currently, scDAPP uses the gene sets from the MSIGDB pathway database, including the Hallmarks pathways, Gene Ontology (GO), KEGG, Reactome, and two transcription factor target databases, the Gene Transcription Regulation Database (GTRD) and Xie et al Nature 2005 database^23–28^. We draw on these databases using the “msigdbr” R package, which also allows flexibility across a range of select species with careful multi-source homology support for gene orthologs^18^.

### Outputs of the scDAPP pipeline

Once completed, scDAPP will save a variety of critical outputs from each stage of the pipeline, including a detailed HTML file (derived from R Markdown; containing extensive visualizations and tables for QC, intermediate results, and final results), text-format results, intermediate data files, and R objects (**Figure 3**). The HTML report file summarizes all steps of the pipeline and all processing and results, including quality control information, data behind threshold selections, UMAPs for clustering results, plots for marker genes, and cross-group analysis. Relevant codes are embedded and can be viewed in this report file.

**Figure 3.**
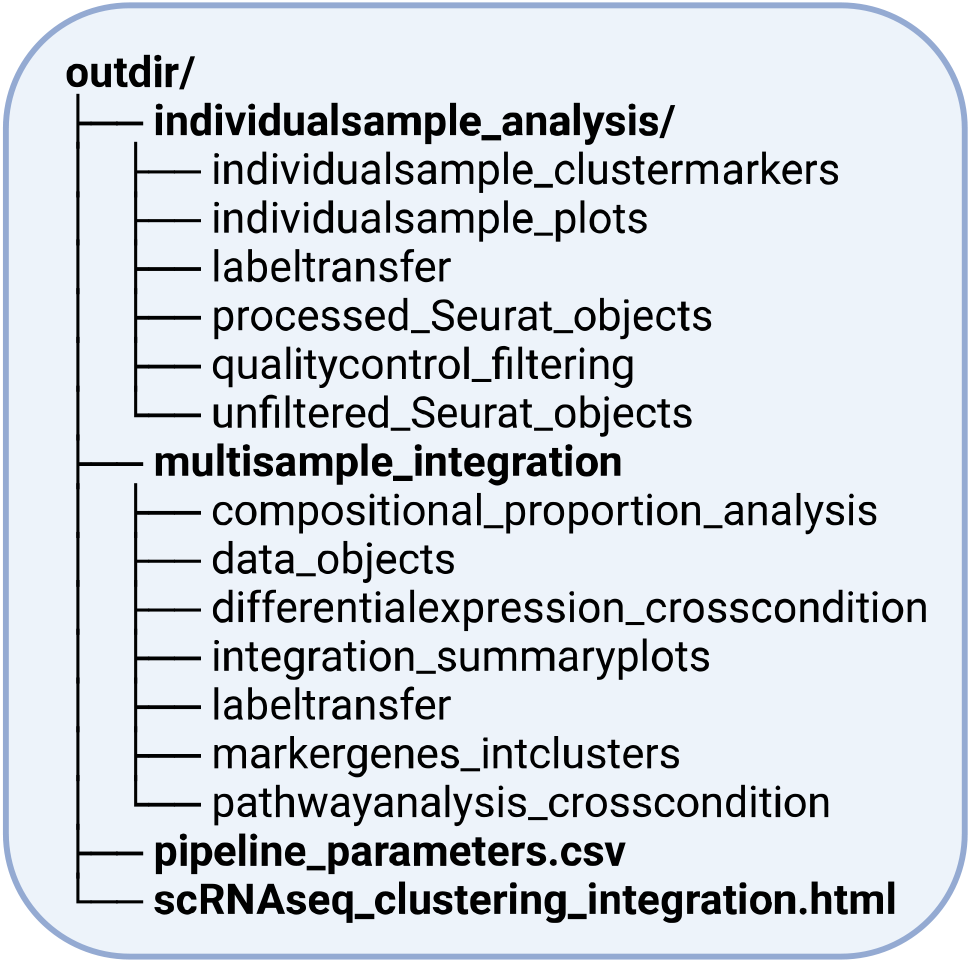
Outputs for scDAPP: result file structures.

Key result tables are saved as .csv files, including results for marker genes, differential expression, differential abundance, and pathway analysis. High-resolution plots are also stored as .pdf files for each step of the pipeline. All these are included to make it transparent for users to track and understand the nuance and decisions in each step of the scRNA-seq analysis. The information is also valuable for users to adjust the parameters to re-run scDAPP until they are satisfied. In this sense, scDAPP is an excellent education tool for learning scRNA-seq analysis. Additionally, relevant data objects are exported and saved, including clustered Seurat objects for each individual sample, a RISC object for the integrated data, and a Seurat object converted from the integrated data in the RISC format. Users with greater bioinformatics expertise can use them to seamlessly conduct further analyses using their own established workflows. Related to this, scDAPP has some utility R scripts for downstream analysis, such as a wrapper function for the recently described aPEAR algorithm for clustering enriched functions from GSEA^29^. Another downstream application includes preparation of interactive web applications. For this purpose, we provide a short vignette linking the output of scDAPP with the ShinyCell package, which is a user-friendly tool to easily export scRNA-seq objects to a web-based Shiny app^30^.

## Results

We have applied scDAPP to many scRNA-seq and snRNA-seq data in our own studies. To demonstrate its performance, we included two instances below.

### Reanalysis of COVID-19 Blood Atlas dataset with scDAPP revealed high concordance with published findings

We applied scDAPP to an atlas dataset of human blood cells from COVID-19 patients and healthy controls^31^. We selected three patient samples each from mild COVID, critical COVID, and controls and ran scDAPP in pseudobulk mode to account for replicates. The input contained 9 samples (3×3) and a total of 83,356 cells. We ran scDAPP on a SLURM-based high performance cluster (HPC) with an allocation of 100GB memory and 9 CPUs, and completed in ∼14 hours (9 x 14 = 126 CPU hours). The main HTML report is in a supplemental data file online (https://github.com/bioinfoDZ/scDAPP/CovidReport_scRNAseq_clustering_integration.html).

The run also used the label transfer option, taking an independent healthy control sample as the reference for cell annotation (provided by the original authors). Overall, our results are in strong concordance with the original publication, for example, increased abundance of plasmablast (PB) and B cells, depletion of NK cells, and overexpression of interferon responses in many cell types in critical COVID [**Figure S1**].

### Reanalysis of mouse developmental scRNA-seq data with scDAPP recapitulated published results

We next applied scDAPP to a scRNA-seq dataset from mouse neural crest cells (NCCs), collected for studying the congenital heart defects observed in 22q11.2 deletion syndrome ^32^. The data were from heart tissues at embryonic day (E)10.5 of control embryos or embryos with *Tbx1* knockout (Wnt1-Cre;Tbx1^-/-^), a gene in the 22q11.2 region and required for cardiac development^33–35^. The input contained four E10.5 samples (2 *Tbx1* wild-type and 2 knockout) and a total of 39,401 cells. We ran scDAPP with standard parameters except for the integrated clustering resolution (“res_int”; set to 1) to obtain clusters closely matching the previous publication, on a SLURM-based HPC with 50 GB memory allocation and 4 CPUs for ∼12 hours. The RPCI reference sample automatically chosen by scDAPP was “Control2”, the same one selected manually in the paper [**Figure S2A**]. Taken the authors’ cell type annotation and markers from an E9.5 sample in the same study for label transfer, scDAPP was able to accurately assign cell types [**Figure S2B;S2C**]. A close examination of the cluster relationship revealed differences, but most cells were clustered similarly by types in scDAPP and previous report [Figure **S2D**], suggesting that computational choice could make subtle difference. The cell composition change between the controls and *Tbx1* null embryos was also reproduced (difference in the craniofacial and outflow tract NCCs, corresponding to the scDAPP clusters 2 and 6) **[Figure S2E]**. Finally, we compared the DEGs in the cardiac progenitor NCCs and found a large agreement between scDAPP and previous results, e.g., upregulation of *Msx2* and *Bambi* with loss of *Tbx1* [**Figure S2F**].

## Discussion

Here we describe scDAPP, a pipeline for single-cell RNA-seq comparative analysis designed with scalability, systematization, and ease-of-use in mind. Whether it is used for a quick end-to-end bioinformatics analysis of the data or recurrent processing of the same data with different options, a main goal is always to help users visualize the granularity of sc/nRNA-seq analysis and achieve a quick transition from raw data to biological interpretation, while learning and exploring analytic parameters and options along the way. Based on our experience, we believe that the visualizations in the HTML output are extremely valuable for users, to examine data distribution, cluster marker specificity, cluster relationship, cell population shifts, biological relevant pathways, and so on.

Importantly, scDAPP places a strong emphasis on full utilization of replicates for both differential gene expression analysis and differential cell composition analysis. The usage of replicates has repeatedly been demonstrated as a critical factor for specificity in single-cell transcriptomic comparative analysis. To our knowledge, scDAPP is the first such end-to-end pipeline to explicitly feature this replicate-aware approach. We should note that there are other computational methods that use replicates but not pseudobulking. Additionally, this pipeline is the first to implement the highly accurate integration software RISC for batch correction and integrated clustering.

As the scDAPP is a pipeline, we have not systematically benchmarked all the software by ourselves and rather have taken the recommendations from the community. The modular design, however, provides sufficient flexibility for including additional software in the future. For example, in addition to Propeller, modified Dirichlet models were shown to perform very well for differential cluster abundance analysis^3^. We may consider it as a second software for cell composition analysis. Additionally, new methods based on combinatorial indexing are now capable of producing datasets with hundreds of thousands to millions of cells per sample^36^. Such technological advancements represent a paradigm shift for the single-cell field in general and may require implementation of specialized tools relying on highly optimized memory storage or combining cells together into metacells. We have not provided such capacities in the scDAPP, but we envision that such approaches may be important to include for future releases.

## Acknowledgement

The authors would like to thank the members in the Zheng’s lab for testing the software and providing valuable suggestions.

