## Supplemental Figures 1 and 2 for "scDAPP: a comprehensive single-cell transcriptomics analysis pipeline optimized for cross-group comparison"

**Supplemental Figure 1. Reanalysis of blood scRNA-seq data from COVID-19 patients and controls with scDAPP**. A) Integrated UMAPs colored by clusters and separated by patient groups. B) Table showing the label transfer result using a healthy non-COVID-19 sample from the same study (not included in the clustering and other re-analysis). MNP = "Mononuclear phagocytes" (monocytes / macrophages), PB = "plasmablast", NK = "Natural Killer cells", "nan" indicates un-annotated cell types from the original published annotations. C) Heatmap showing the expression of markers computed for the clusters in A. D) Heatmap table showing the cell composition change between critical COVID-19 patient and healthy control samples. E) Differential pathway analysis showing gene sets upregulated in critical COVID-19 patients versus healthy samples. ** All these figures were taken from the scDAPP output directly.

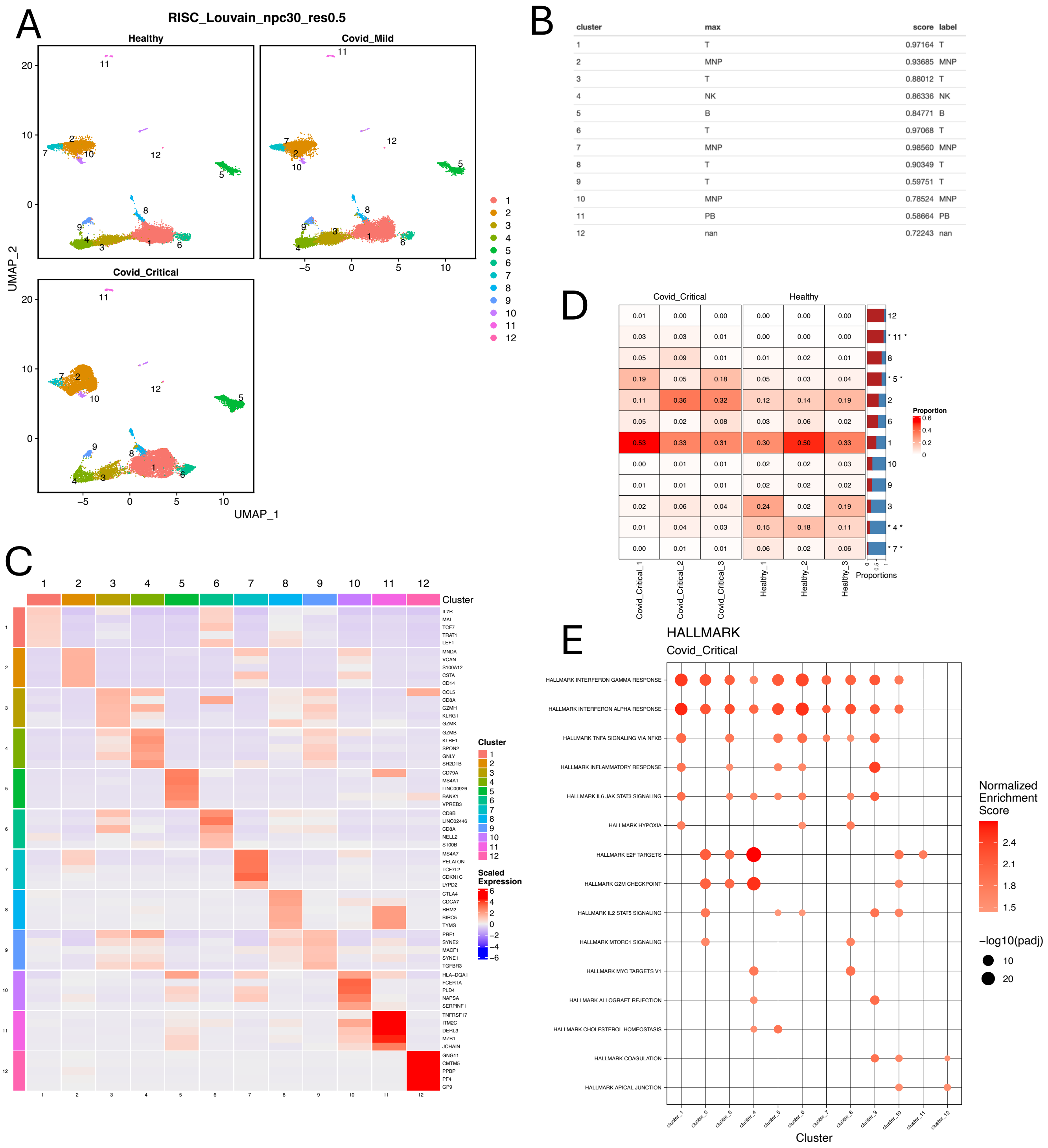

**Supplemental Figure 2. Reanalysis of E10.5 Neural Crest Cell (NCC) scRNA-seq data with scDAPP.** A) Table showing the scDAPP-computed RISC reference scores for the four samples, indicating that the sample Control2 had the highest score and was selected as the reference for RISC integration, consistent with what was selected manually in the original analysis. B) Table showing the cluster annotation from label transfer using an E9.5 NCC dataset, yielding the same cluster annotation in the original publication. C) UMAP plot of the integrated samples colored by clusters. D) Table showing the relationship between clusters in published versus scDAPP re-analysis. Note that clusters 6 and 17 were not included in the original study, indicating different thresholds for cell filtering. E). Table for cell compositional analysis showing significant decreases in cell number with *Tbx1* KO in the clusters 2 and 6, consistent with the original report. F) Violin plots showing the differential expression of four key genes highlighted in the paper for cardiac progenitor neural crest cells (cluster 3 in scDAPP). Panels A, B, C, and E were taken directly from the scDAPP output.

A

| Code | Set | RefScore | Max |
|---|---|---|---|
| Control1 | Set-1 | 23.87617 | |
| Control2 | Set-2 | 35.46275 | * |
| KO1 | Set-3 | 25.04691 | |
| KO2 | Set-4 | 24.44501 | |

B

| cluster | max | score | label |
|---|---|---|---|
| 1 | NT/Neural1 | 0.68682 | NT/Neural1 |
| 2 | NCCs-DistalPA1-2 | 0.63041 | NCCs-DistalPA1-2 |
| 3 | NCCs-ProximalPA1 | 0.48151 | NCCs-ProximalPA1_putative |
| 4 | NT/Neural1 | 0.73458 | NT/Neural1 |
| 5 | NCCs-EarlyMigrating-PA1 | 0.52825 | NCCs-EarlyMigrating-PA1 |
| 6 | NCCs-ProximalPA1 | 0.60004 | NCCs-ProximalPA1 |
| 7 | NCCs-ProximalPA1 | 0.63901 | NCCs-ProximalPA1 |
| 8 | NT/Neural2 | 0.89519 | NT/Neural2 |
| 9 | NT/Neural5 | 0.92197 | NT/Neural5 |
| 10 | NT/Neural3 | 0.90672 | NT/Neural3 |
| 11 | NT/Neural5 | 0.93957 | NT/Neural5 |
| 12 | SM-CNCCs | 0.42429 | SM-CNCCs_putative |
| 13 | Facial-NCCs | 0.30365 | Facial-NCCs_putative |
| 14 | NT/Neural2 | 0.51081 | NT/Neural2 |
| 15 | Neuronal | 0.86334 | Neuronal |
| 16 | NT/Neural5 | 0.60665 | NT/Neural5 |
| 17 | NCCs-ProximalPA1 | 0.43511 | NCCs-ProximalPA1_putative |

C

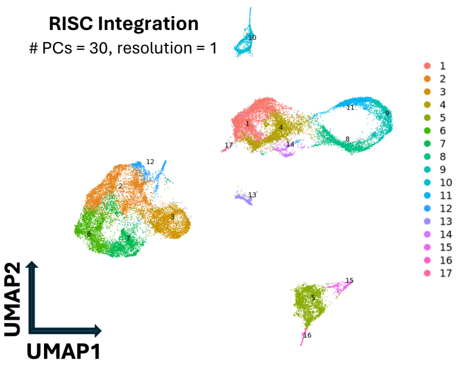


D

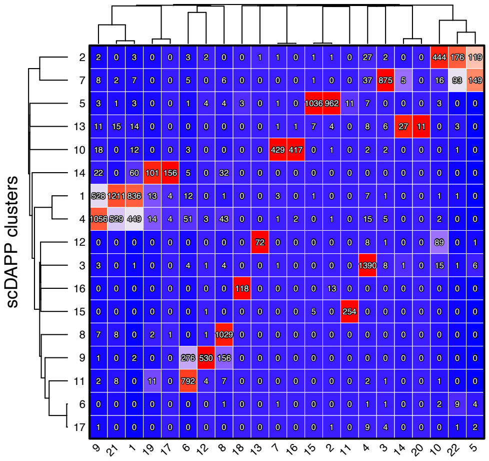


E

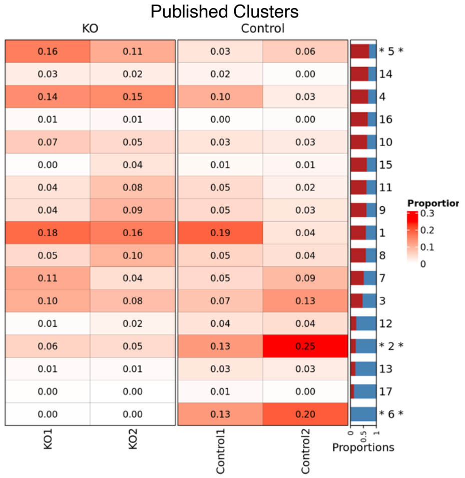


F

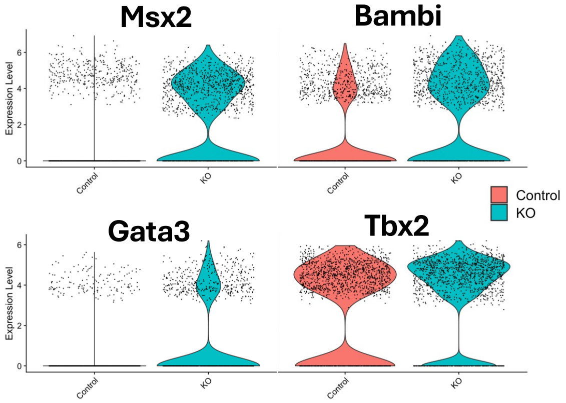
